# Environmentally-induced mdig is a major contributor to the severity of COVID-19 through fostering expression of SARS-CoV-2 receptor NRPs and glycan metabolism

**DOI:** 10.1101/2021.01.31.429010

**Authors:** Qian Zhang, Chitra Thakur, Yao Fu, Zhuoyue Bi, Yiran Qiu, Priya Wadgaonkar, Liping Xu, Bandar Almutairy, Wenxuan Zhang, Paul Stemmer, Fei Chen

## Abstract

The novel β-coronavirus, SARS-CoV-2, the causative agent of coronavirus disease 2019 (COVID-19), has infected more than 101 million people and resulted in 2.2 million death worldwide. Recent epidemiological studies suggested that some environmental factors, such as air pollution, might be the important contributors to the mortality of COVID-19. However, how environmental exposure enhances the severity of COVID-19 remains to be fully understood. In the present report, we provide evidence showing that mdig, a previously reported environmentally-induced oncogene that antagonizes repressive trimethylation of histone proteins, is a master regulator for SARS-CoV-2 receptors neuropilin-1 (NRP1) and NRP2, cathepsins, glycan metabolism and inflammation, key determinants for viral infection and cytokine storm of the patients. Depletion of mdig in bronchial epithelial cells by CRISPR-Cas-9 gene editing resulted in a decreased expression of NRP1, NRP2, cathepsins, and genes involved in protein glycosylation and inflammation, largely due to a substantial enrichment of lysine 9 and/or lysine 27 trimethylation of histone H3 (H3K9me3/H3K27me3) on these genes as determined by ChIP-seq. These data, accordingly, suggest that mdig is a key mediator for the severity of COVID-19 in response to environmental exposure and targeting mdig may be one of the effective strategies in ameliorating the symptom and reducing the mortality of COVID-19.

## Introduction

The global outbreak of COVID-19 caused by the newly identified SARS-CoV-2 virus, is having a devastating impact on both public health and social economics. According to the World Health Organization (WHO), as of January 30, 2021, there had been more than 101 million confirmed cases of COVID-19 worldwide, leading to almost 2.2 million deaths. The SARS-CoV-2 virus has a large number of trimeric spikes (S) on its surface. Each S protein has two subdomains, S1 and S2. Upon direct binding of S1 to the ACE2 on the surface of the host cells, membrane-associated proteases of the host cells, including furin, TMPRSS2, and others, cleave S proteins at the RRAR/S motif located at the S1-S2 boundary, or the motif of PSKR/S, the so-called S2’ site, in S2 region, followed by S2-mediated membrane fusion ^1,2^. In addition to ACE2, the C-terminal RRAR motif of the S1 protein from the cleaved S protein may bind to cell surface neuropilin-1 (NRP1) or NRP2 for sufficient viral entry into the host cells ^3–5^. After membrane fusion, the virus is shuttled in the cytoplasm through the endosomes and lysosomes, during which several cathepsin family proteases cleave S proteins further and other viral proteins, leading to the release of the RNA genome of SARS-CoV-2 into the cytoplasm ^2^. Indeed, some inhibitors targeting endosome, lysosome or cathepsin, such as ammonia chloride, bafilomycin A and teicoplanin, had been shown to be effective in blocking viral entry in cell-based experiments ^2,6^. The viral RNA genome can hijack the protein translational machinery of the host cells for the expression of viral polyproteins that can be processed to generate replicase, proteases, transcriptases, and other viral structural proteins for assembly of the new virion. Some of these newly generated viral proteins can antagonize the anti-viral responses of the interferon signaling, whereas others are highly capable of activating the inflammasome for cytokine storm and pyroptosis of the host cells in the lung and other organs, leading to an increased likelihood of fatality of the COVID-19 patients ^7^.

Both S protein and ACE2 are known to be extensively glycosylated through covalently linked oligosaccharides (glycans). It is believed that glycosylation may enhance the infectivity of the virus. Many glycoconjugates on the surface of the host cells are also modified by sialic acid, a specific form of glycosylation--sialylation, which may facilitate membrane fusion of the virus and cells. It is also very likely that alteration in ACE2 and membrane glycan profile may determine the inter-individual variation on the outcomes of viral infection ^8^.

Human lung is the first line of attack by the SARS-CoV-2 virus. The fatality of COVID-19 has been largely attributed to the massive alveolar damage and progressive respiratory failure resulted from cellular fibromyxoid exudates, formation of hyaline (hyaluronan) membrane, pulmonary oedema, inflammation, and interstitial fibrosis ^9–11^. Most environmental risk factors, such as air pollution and PM2.5, can induce chronic inflammation and make the lung more vulnerable to viral attacks. Indeed, a recent nationwide epidemiological study in the US had shown that an increase of only 1 *μ*g/m^3^ in PM2.5 is associated with an 8% increase in the COVID-19 death rate, clearly indicating that environmental factors may contribute to the severity of the disease outcomes among individuals with COVID-19 ^12^. This notion is complementarily supported by the fact that the areas, such as Lombardy, Emilia-Romanga, Piemonte, and Veneto of Italy, and Madrid of Spain, with worse air quality due to higher level of nitrogen dioxide (NO2), are the hardest-hit areas with deaths from COVID-19 ^13^.

The mineral dust-induced gene mdig was first identified in alveolar macrophages from people with chronic lung diseases associated with occupational exposure ^14^, and additional studies concluded that mdig can be induced by a number of environmental hazards, such as silica particles, arsenic, tobacco smoke, and PM2.5 ^14^. This gene was independently discovered as myc-induced nuclear antigen (mina53) and nucleolar protein NO52, respectively. Functional tests suggested that mdig may have hydroxylase activity on ribosomal protein L27a, and accordingly, an alternative name, Riox2, was given ^14^. The mdig protein contains a conserved JmjC domain that was considered as a signature motif of the histone demethylase family members. In human bronchial epithelial cells, lung cancer cell line A549, and breast cancer cell line MDA-MB-231 cells, knockout of mdig gene by CRISPR-Cas9 gene editing, resulted in a pronounced enrichment of histone H3 lysine9 trimethylation (H3K9me3) as well as H3K27me3 and H4K20me3 in ChIP-seq analysis, esp. in the gene loci encoding proteins in inflammation and fibrosis, such H19, TGFβ signaling, collagens, and cell adhesion molecules ^15^. Data from mice with heterozygotic deletion of mdig gene indicated that mdig is a master regulator of inflammation and tissue fibrosis ^15,16^. In the present report, we further demonstrated that mdig may exacerbate the severity of COVID-19 in response to environmental exposure.

## Results

### Depletion of mdig prevents cleavage of SARS-CoV-2 spike protein

We had used non-cancerous bronchial epithelial cell line BEAS-2B to establish mdig knockout (KO) cells through CRISPR-Cas9 gene editing, and used the cells subjected to gene editing but without mdig depletion as wild type (WT) cells ^15^. By transfection of the WT and KO cells with an expression vector for the full-length spike (S) protein of SARS-CoV-2, a detectable decrease in S protein cleavage was noted in the mdig KO cells (Fig. 1A, top panel). The molecular weight (MW) of the unprocessed full-length S protein is around 200 kDa (green arrow), and the cleaved product is about 100-110 kDa (red arrow). To determine whether the decreased cleavage of S protein in mdig KO cells is a result of the diminished expression of proteases responsible for S protein cleavage, we compared the protein levels of cathepsin D (CTSD), transmembrane serine protease 2 (TMPRSS2), and furin between WT and mdig KO cells. There is no measurable difference in the levels of TMPRSS2 and furin. However, a substantial decrease of CTSD, both the 46 kDa precursor and 28 kDa mature form, was observed in the mdig KO cells (Fig. 1A). In another cell line, the triple-negative breast cancer cell line MDA-MB-231 cell, we noted that CTSD is the most diminished protein when mdig was knocked out in this cell line (data not shown).

**Fig. 1.**
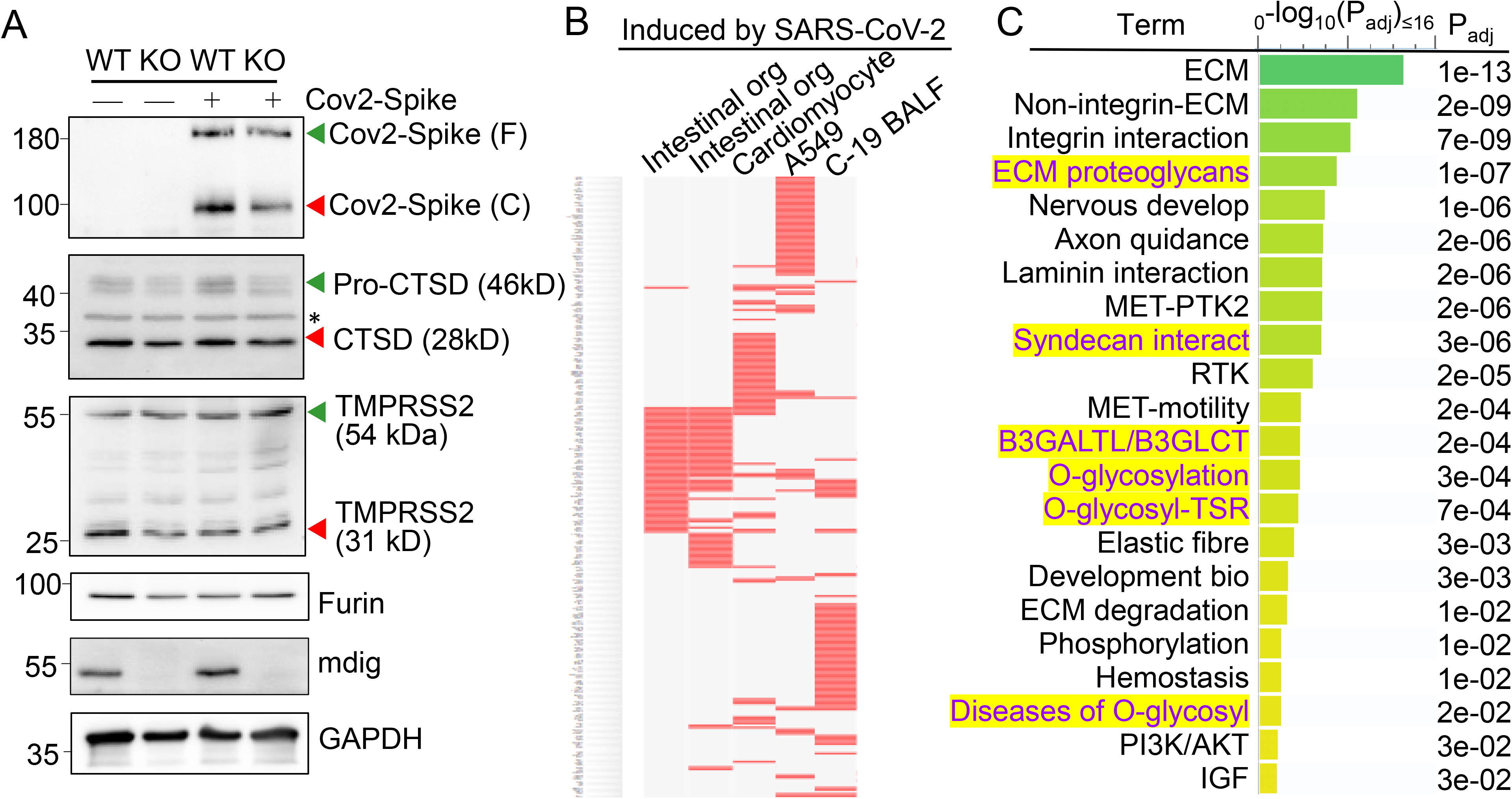
Knockout of mdig diminishes the expression of genes for SARS-CoV-2 infectivity. **A.** Western blotting shows decreased cleavage of the transfected SARS-CoV-2 S protein and the expression of CTSD in BEAS-2B cells. **B.** Down-regulated genes in the mdig KO cells as determined by RNA-seq are highly represented for those gene induced by SARS-CoV-2 in intestinal organoids (GSE149312), cardiomyocytes (GSE150392), A549 cells (GSE147507), and bronchoalveolar lavage fluid (BALF) of COVID-19 patients. **C.** Reactome pathway assay shows the down-regulated genes in mdig KO cells as determined by RNA-seq are mostly in the pathways of extracellular matrix (ECM) regulation and glycan metabolism (highlighted in yellow).

To additionally investigate the effect of mdig knockout on the genes linked to SARS-CoV-2 infection and pathology, we performed gene expression profiling between WT and mdig KO cells by two independent RNA-seq analyses. The differentially expressed genes identified in both sets of RNA-seq data were then subjected to gene set enrichment analysis with the Enrichr web-based application. This analysis showed that the down-regulated genes in the mdig KO cells were over-presented in the gene sets induced by SARS-CoV-2 in intestinal organoids (GSE149312), cardiomyocytes (GSE150392), A549 cells (GSE147507), and bronchoalveolar lavage fluid (BALF) of COVID-19 patients (Fig. 1B), suggesting that mdig enhances expression of genes involved in either SARS-CoV-2 infection or the pathogenesis of COVID-19, To further understand the biological functions of the down-regulated genes identified in RNA-seq in mdig KO cells, we applied enrichment assay of the Reactome. In this analysis, we found that the top Reactome pathways of these down-regulated genes in mdig KO cells are related to organization, interaction and regulation of extracellular matrix (ECM) (Fig. 1C). Unexpectedly, six of the twenty-two top-ranked Reactome pathways are centered on protein glycosylation or glycan metabolism, indicating that in addition to ECM, mdig is critical for the post-translational modification of proteins by carbohydrate moieties.

### Involvement of mdig in antagonizing H3K9me3 and/or H3K27me3

A previous study showed that knockout of mdig in BEAS-2B cells caused significant enrichments of H3K9me3, H3K27me3, and H4K20me3 on the merged peak regions as determined by ChIP-seq ^15^. Re-analysis of the ChIP-seq data from the WT and mdig KO cells by pairwise Pearson correlation analyses, through plotting the tag numbers of H3K4me3, H3K9me3 and H3K27me3 in the promoter region in mdig KO cells against the corresponding tag numbers in WT cells, revealed a significant enhancement of H3K9me3 and H3K27me3 in gene promoters in mdig KO cells (Fig. 2A). Clustered heatmaps as depicted in Fig. 2B suggested that the enrichment of H3K9me3 and H3K27me3 were drastically enhanced in clusters 2 and 3 of the mdig KO cells, while little to no changes in the enrichment of H3K9me3 and H3K27me3 were observed in clusters 1, 4 and 5. The enhanced H3K9me3 in mdig KO cells was also supported by the significantly increased number of H3K9me3 peaks in ChIP-seq. In WT cells, the normalized number of H3K9me3 peaks is 12,094, whereas in KO cells, this number is 32,264 (data not shown). To determine which sets of genes are enriched with H3K9me3 in mdig KO cells, we conducted Reactome pathway assay for those H3K9me3-enriched genes. It is interesting to note that the top-ranked pathways are mostly associated with protein glycosylation and glycan regulation (Fig. 2C), which is highly correlated to the pathways of the down-regulated genes in RNA-seq of the mdig KO cells (Fig. 1C). These data, thus, provide an unequivocal evidence indicating that mdig fosters the expression of genes in glycosylation and glycan metabolism through antagonizing the repressive histone methylation marker H3K9me3 as well as H3K27me3.

**Fig. 2.**
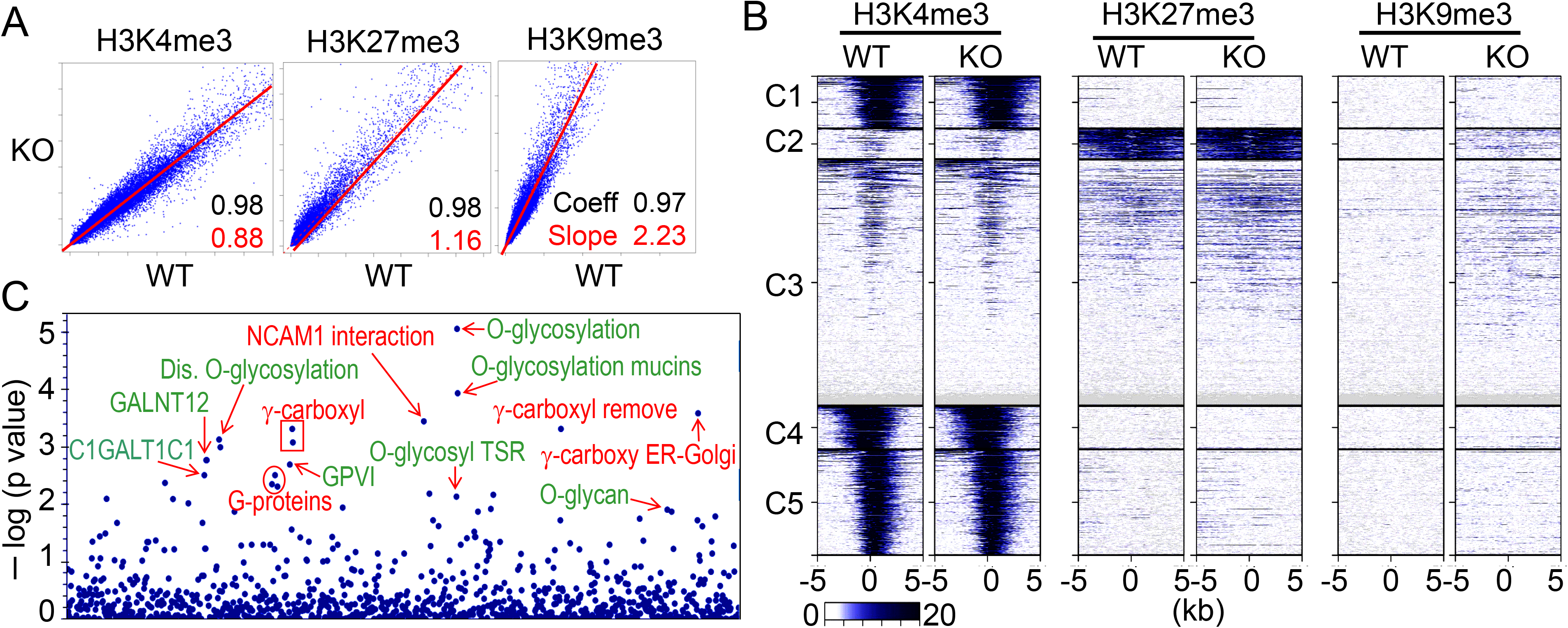
ChIP-seq data revealed that mdig is antagonistic to H3K27me3 and H3K9me3. **A.** Pairwise Pearson correlation analyses were made through plotting the tag numbers of H3K4me3, H3K9me3 and H3K27me3 in the promoter region in mdig KO cells against the corresponding tag numbers in WT cells. **B.** Clustered heatmaps of ChIP-seq for H3K4me3, H3K27me3 and H3K9me3 in the promoter regions of WT cells and mdig KO cells. **C.** Reactome pathway analysis of the genes showed significantly enhanced enrichment of H3K9me3 in mdig KO cells. The top-ranked pathways in glycan metabolism are marked in green color.

### Receptors and proteases required for SARS-CoV-2 infection are regulatory targets of mdig

Considering the fact that knockout of mdig reduced cleavage of SARS-CoV-2 spike (S) protein (Fig.1A), we narrowed down our analysis of ChIP-seq and RNA-seq data to several key proteins important for S protein binding, intracellular vesicle trafficking and fusion. Upon close examination, we found that genes involved in key aspects of SARS-CoV-2 entry into the host cells, including NRP1, NRP2, DPP4, TMPRSS15, RAB6B, IQGAP2, and others, are enriched with the repressive histone trimethylation marker, H3K9me3 or H3K27me3, along with a decreased expression of these genes as determined by RNA-seq, in mdig KO cells (Fig. 3A). NRP1 and NRP2 are transmembrane proteins serving as receptors for growth factors and semaphorins. Recent reports suggest that NRP1 or NRP2 can function as SARS-CoV-2 receptor through binding of the C-terminal RRAR motif of the furin-cleaved S protein, followed by internalization of the virus via micropinocytosis ^3,4^. Knockout of mdig raised H3K9me3 in the up- and down-stream of NRP1 gene, and H3K27me3 and H3K9me3 in the gene body and down-stream of NRP2 gene, which are consistent with the diminished gene expression as determined by RNA-seq (Fig. 3A). Meanwhile, a drastic reduction of H3K4me3, an active transcription marker, was observed on both NRP1 and NRP2 genes in mdig KO cells, which further enforced the transcriptional suppression of NRP1 and NRP2 by an increased level of H3K9me3 and/or H3K27me3. The role of mdig on NRP1 was also confirmed in both MDA-MB-231 breast cancer cells and A549 lung cancer cells. Knockout of mdig enhanced enrichment of H3K9me3 in the upstream of NRP1 gene, and H4K20me3, another repressive histone methylation marker, in the gene body and upstream of the NRP1 gene in MDA-MB-231 cells (sFig. 1), which correlated to a 2.5-fold decrease of the NRP1 protein in the KO cells as determined by quantitative proteomics (sFig.2). In A549 cells, knockout of mdig resulted in an enhancement of H4K20me3 in the down-stream of the NRP1 gene (sFig. 3). The mdig KO cells also exhibited reduced expression of DPP4 and TMPRSS15 due to intensified enrichment of H3K9me3 on these genes. DPP4 serves as a receptor for MERS-CoV and possibly for SARS-CoV-2 too ^17^. Inhibitors of DPP4 conferred a 7-fold lower risk of COVID-19 mortality among patients with metabolic diseases ^18^. TMPRSS15 shares high homology in the serine protease domains of TMPRSS2, thus, may participate in the cleavage of the S protein of SARS-CoV-2 during infection. Both RAB6B and IQGAP2 are important regulators for lysosome-phagosome tethering and endocytic vesicle trafficking or fusion ^19^. Thus, the enhanced enrichment of H3K9me3 and/or H3K27me3 on and down-regulation of these genes in mdig KO cells suggest that mdig may facilitate SARS-CoV-2 infection through antagonizing the repressive histone methylation markers H3K9me3 and H3K27me3, leading to an upregulated expression of these genes. This notion is further supported by the decreased expression of the cathepsin (CTS) family members due to enhanced enrichment of H3K9me3 on these genes in mdig KO cells (Fig. 3B). Knockout of mdig caused strong enrichments of H3K27me3 and H3K9me3 in the up-stream of CTSD gene, which is consistent to the diminished CTSD protein in mdig KO cells as shown in Fig. 1A. Increased level of H3K9me3 was also seen on the gene bodies of CTSE, CTSL1, CTSL3P, and CTSL1P8 in mdig KO cells (Fig. 3B). Results from RNA-seq showed that except CTSF, the expression levels of all other members of the CTS family are significantly declined following mdig knockout (Fig. 3C). It had been well-documented that CTS members are the dominating proteases for further degradation of SARS-CoV-2 S proteins in lysosomes ^20,21^, which is essential for the release of the viral RNA genome into the cytoplasm.

**Fig. 3.**
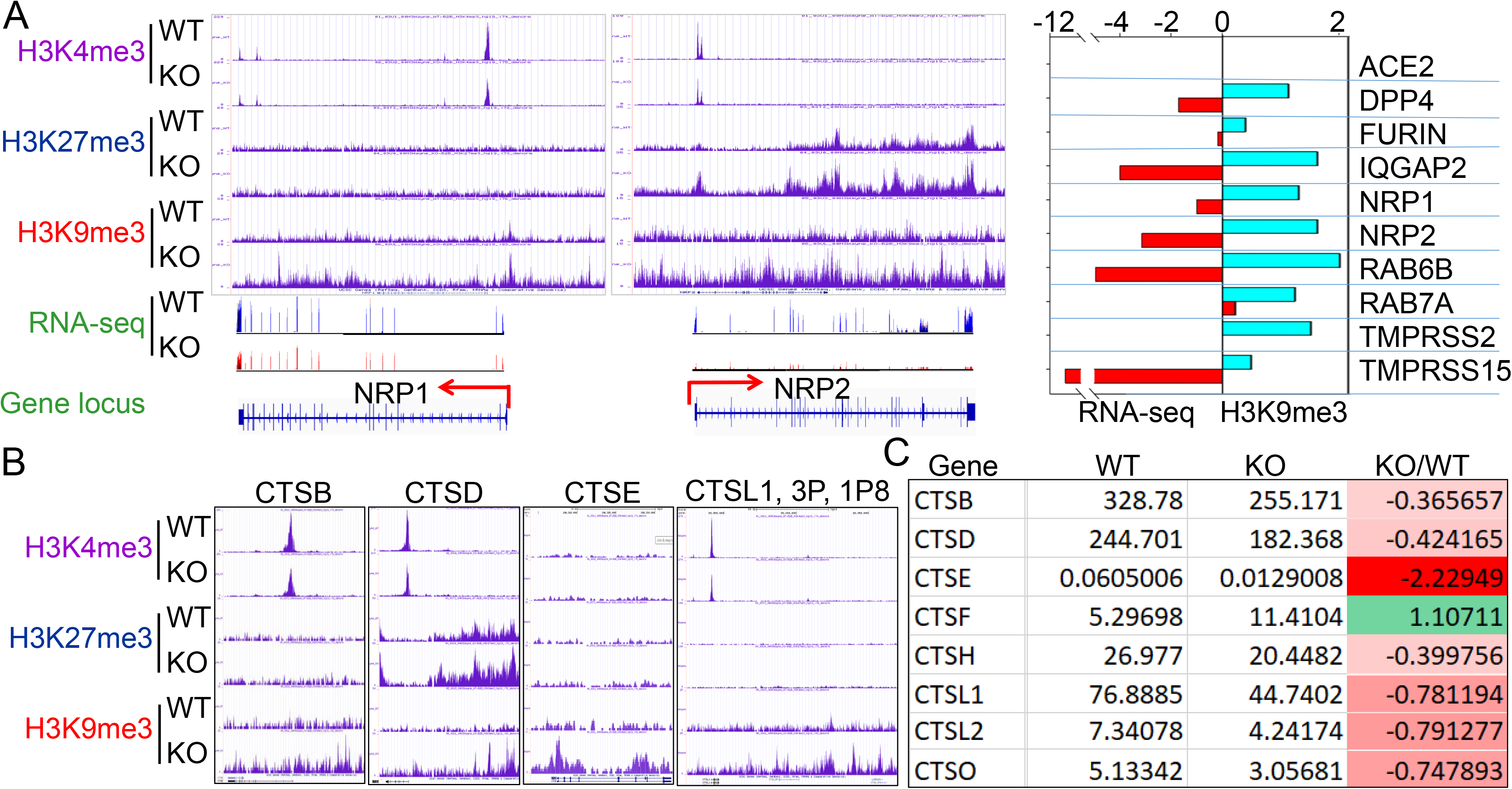
Diminished expression of genes related to SARS-CoV-2 infectivity. **A.** Down-regulation of S protein receptors NRP1 and NRP2 in mdig KO cells. Both ChIP-seq screenshot and RNA-seq spectrum for NRP1 and NRP2 were shown. Right panel summarized the elevated enrichment of H3K9me3 as determined by ChIP-seq and reduced expression in RNA-seq of these indicated genes in mdig KO cells. **B.** Knockout of mdig elevated enrichment of H3K9me3 and/or H3K27me3 on the indicated cathepsin genes that also showed decreased enrichment of the active transcription marker, H3K4me3. **C.** RNA-seq showed down-regulation of these indicated cathepsin genes in the mdig KO cells (except CTSF).

### The expression of the major glycosylation pathway genes is mdig dependent

Glycosylation creates great structural and functional diversity of the target proteins. During SARS-CoV-2 infection, glycosylation of the viral proteins is essential for the assembly of the virion and is able to shield the virus from the host immune response. One of the major glycosylation pathways is the hexosamine biosynthetic pathway (HBP) from glycolytic metabolism of the glucose, which generates uridine diphosphate N-acetylglucosamine (UDP-GlcNAc), the founding molecule for N- and O-glycosylation, sialylation, formation of hyaluronan, and biosynthesis of glycosylposphatidylinosital (GPI) (Fig. 4A). In the mdig KO cells, we observed a strong enrichment of the repressive histone marker, H3K9me3, on the genes in this pathway, such as ST3GAL1 (Fig. 4B) and HAS3 (Fig. 4C). RNA-seq confirmed a decreased expression of these two genes and other genes as indicated in Fig. 4A. Interestingly, most of the GalNAc kinases (GALs) that exhibited increased enrichment of H3K9me3 or H3K27me3 in ChIP-seq, are down-regulated as determined by RNA-seq in the mdig KO cells (Fig. 4D). Accordingly, we believe that in the normal cells, induction of mdig will enforce the overall glycosylation for both viral and host cell proteins.

**Fig. 4.**
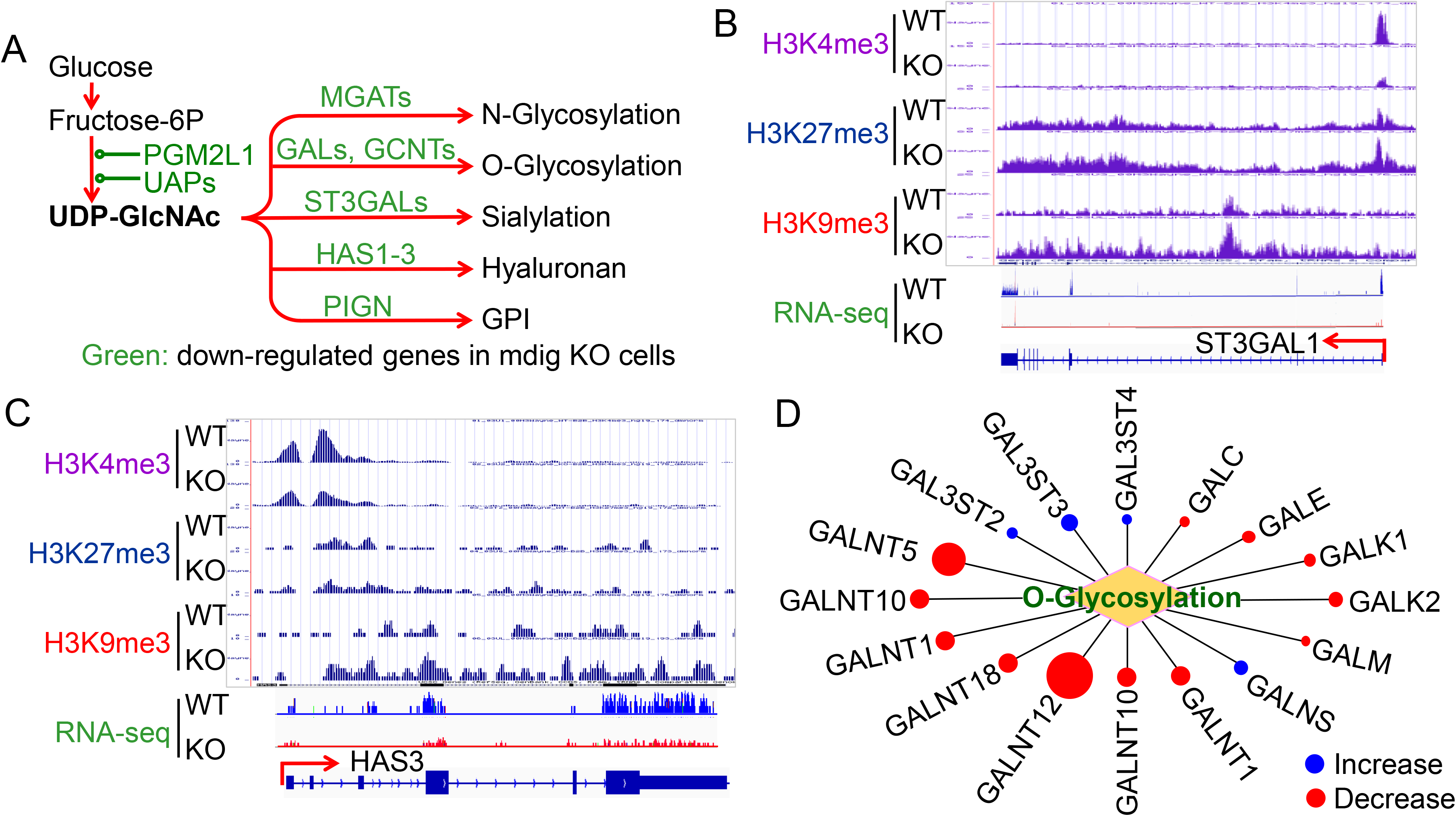
Decreased expression of genes in protein glycosylation. **A.** Diagram shows glucose metabolism and hexosamine biosynthetic pathway linked to glycan metabolism and glycosylation. The down-regulated genes in mdig KO cells are marked in green. **B and C.** Screenshot of ChIP-seq and RNA-seq for ST3GAL1 that catalyzes sialylation and HAS3 that catalyzes the generation of hyaluronan, respectively. Knockout of mdig enriched H3K9me3 and decreased H3K4me3, leading to the diminishment of the ST3GAL1 and HAS3 mRNA in RNA-seq. **D.** Down-regulation of O-glycosylation pathway genes in mdig KO cells. Red circles denote decreased, while blue circles denote an increased expression of the genes in mdig KO cells. The sizes of circles indicate the degree of increase or decrease of these gene expressions in mdig KO cells.

### Mdig is a master regulator of inflammation

Inflammatory cytokine storm is the most important mortality factor of COVID-19 patients. Our recently published data demonstrated that mdig controls the expression of genes involved in inflammation, including TGFβ signaling, collagens and cell adhesion molecules ^14,15^. In addition, heterzygotic knockout of mdig in mice ameliorated silica-induced lung fibrosis along with a significant decrease of Th17 cell and macrophage infiltration into the lung interstitium ^15,16^. Re-analyzing of the ChIP-seq and RNA-seq data from the WT and mdig KO cells further unraveled that mdig regulates a wide spectrum of inflammatory genes, such as several receptor genes for IL17 and IL1 (Figs. 5A and 5B). Knockout of mdig caused an elevated enrichment of H3K27me3 and H3K9me3, both are repressive markers for gene transcription, on IL17RD. Meanwhile, the expression of some TLR and NLRP members involved in inflammasome-mediated inflammation are also compromised in the mdig KO cells (Figs. 5C and 5D). On the genes encoding prostaglandin E2 receptors 2, 3 and 4, although there was no significant gain of H3K9me3 and H3K27me3, a notable reduction of H3K4me3, an active marker for gene transcription, on these genes was observed in the KO cells (Fig. 5C).

**Fig. 5.**
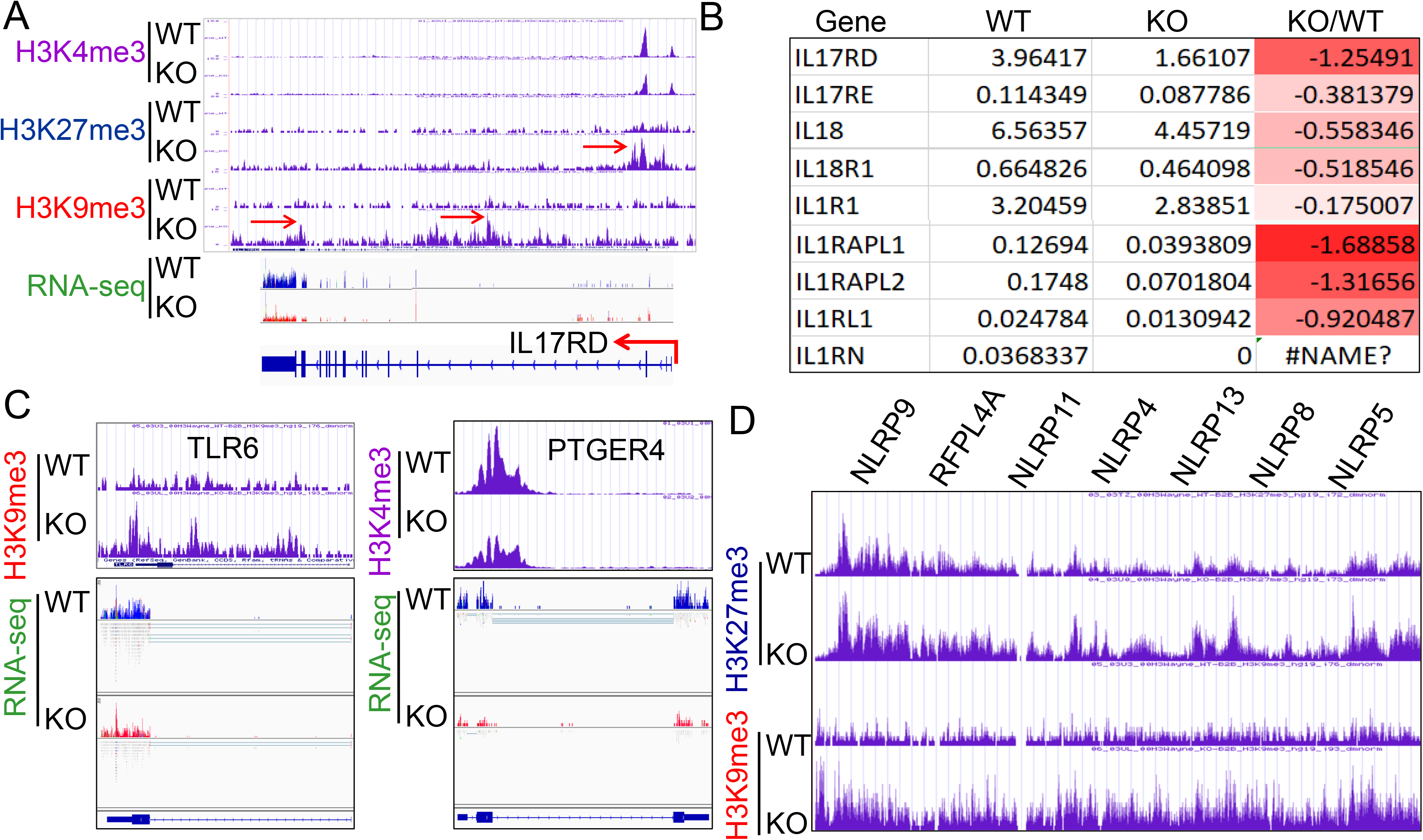
Regulatory role of mdig on inflammation. **A.** ChIP-seq and RNA-seq of IL17RD in both WT and mdig KO cells. Red arrows point to the enhanced peaks of H3K27me3 and H3K9me3 on the IL17RD gene in mdig KO cells. **B.** RNA-seq shows down-regulation of these indicated inflammatory cytokine receptors or their regulatory protein in mdig KO cells. **C.** ChIP-seq shows increased enrichment of H3K9me3 on TLR6 gene and decreased enrichment of H3K4me3 on PTGER4 gene in mdig KO cells, which correlated to the diminished expression of these genes in RNA-seq. **D.** Enrichment of H3K27me3 and H3K9me3 on the NLRP gene loci that encode the key components of inflammasomes.

### Elevated expression of mdig in pulmonary fibrosis

Pulmonary fibrosis is one of the major contributing factors to the acute respiratory distress syndrome (ARDS) of the COVID-19 patients, which is also a common sequela in some COVID-19 survivors ^22,23^. Many fibrotic damages are irreversible and may cause lifelong functional impairment of the lung. The pro-fibrotic role of mdig was first demonstrated in silica-induced lung fibrosis in mice ^14–16^. It is unknown whether mdig is also important in the development of lung fibrosis in humans. To answer this question, we screened mdig expression through immunohistochemistry in tissue microarrays containing 17 normal human lung tissues and 26 cases of fibrotic lung tissues collected from patients with cancer, chronic bronchitis or chronic pneumonia. In agreement with earlier reports that mdig was barely detectable in normal lung tissue ^24,25^, in this analysis, only 24% of normal lung tissues showed some faint staining of mdig. In contrast, the majority of fibrotic lung tissues, 85%, exhibited a strong signal of mdig expression (Fig. 6). Although we cannot distinguish the question of whether mdig promoted fibrosis or fibrosis caused a higher expression of mdig in human lung tissues, based on previous mouse model with mdig knockout and the fact that mdig is a master regulator of inflammation ^14–16^, we believe that mdig is a critical driving factor for the development of lung fibrosis associated with some disease conditions, such as COVID-19, chronic bronchitis, pneumonia, and cancer.

**Fig. 6.**
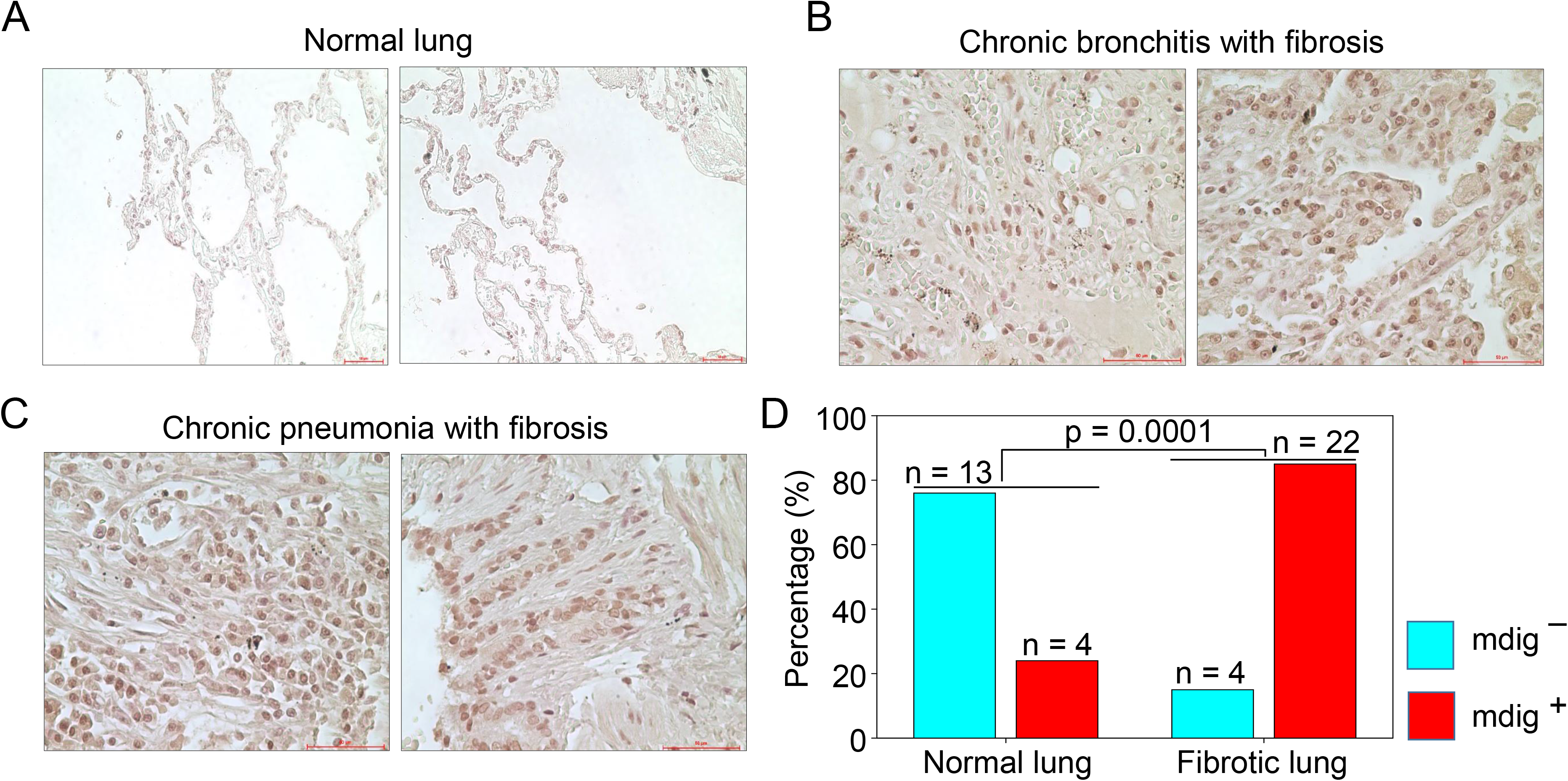
Immunohistochemical staining of mdig in normal and fibrotic human lung tissues. **A.** Staining of mdig in representative normal human lung tissues. A total of 17 normal lung tissue samples were investigated. **B.** Staining of mdig in fibrotic lung tissues from patients with chronic bronchitis. **C.** Staining of mdig in fibrotic lung tissues from patients with chronic pneumonia. **D.** Summary of mdig positive rates in normal and fibrotic human lung tissues.

## Discussion

The COVID-19 pandemic due to SARS-CoV-2 infection has many destructive effects on public health and social-economics on a global scale. Despite a large amount of molecular detail is known about the virology of SARS-CoV-2 and the pathology of COVID-19, little is currently known about why some COVID-19 patients showed mild or no symptoms but others, even who are at young ages and have no pre-existing disease conditions, have severe or fatal outcomes. A number of studies had demonstrated that several environmental factors, including air pollution, temperature, humidity, wind speed, etc. had a significant influence on the mortality of COVID-19 ^12,13,26^. However, to the best of our knowledge, no detailed studies have investigated how these suspected environmental factors intensify the severity or mortality of COVID-19. In this report, we provided evidence suggesting that mdig, a potential oncogenic gene induced by several common environmental hazards ^14,27^, may augment the infectivity of SARS-CoV-2 and the severity of COVID-19 through upregulating S protein receptors NRP1 and NRP2, several proteases and the genes in the pathways of glycan metabolism and inflammation.

It has been a general assumption that ACE2 is abundantly expressed in bronchial epithelial cells and other lung cells, and serves as the main receptor for the S protein of SARS-CoV-2 ^28^. It was beyond our expectation that ACE2 was barely detected by RNA-seq in both WT and mdig KO BEAS-2B cells, a cell line derived from non-cancerous bronchial epithelial cells. Although we cannot rule out the possibility of genetic or epigenetic variations during the establishment and passage of the cell lines, there is an evidence showing very low protein levels of ACE2 in respiratory epithelial cells ^29^. Accordingly, it is very likely that NRP1 and NRP2 are critically required for efficient viral entry into the host cells in the lung. The dependence of mdig on the expression of NRP1 and NRP2 was also observed in two additional cell lines, MDA-MB-231 (sFig.1 and sFig. 2) and A549 (sFig. 3). Thus, it is plausible to speculate that mdig dependent expression of NRP1 and NRP2 can enhance the infectivity of SARS-CoV-2, leading to the worsening effect of environmental exposure on the severity of COVID-19.

Sequential cleavage of the S protein at S1-S2 boundary and S2’ site by host proteases is a prerequired step for conformational changes and membrane fusion of the S protein, followed by viral internalization and intracellular trafficking within endosomes and lysosomes. Inside the lysosomes, the S proteins on the surface of SARS-CoV-2 will be subjected to additional cleavages by cathepsins, primarily by CTSL and CTSB. A number of studies concluded that inhibitors that target lysosome or the lysosomal proteases, cathepsins, can prevent the release of the viral RNA genome into the host cytoplasm and viral entry into the target cells ^30^. Decreased expression of CTSB, CTSD, CTSE, and CTSL members in the mdig KO cells suggests that mdig bolsters the expression of proteases required for additional cleavage of the S protein, which provides a permissive environment for SARS-CoV-2 propagation.

Both S protein of SARS-CoV-2 and ACE2 are highly glycosylated ^31^. Certainly, glycosylation changes the antigenicity of S protein and its affinity towards cellular receptors. Meanwhile, membrane fusion of SARS-CoV-2 with the host cells is strengthened by some types of glycosylation of the cell surface proteins, such as sialylation ^32^. More importantly, in addition to intensify the infectivity of the SARS-CoV-2, amplified glycosylation resulted from the excessive generation of hyaluronan that forms liquid jelly-like structure in both alveolar space and the interstitial area is fatal for COVID-19 patients ^33^. Inflammatory cytokine storm is one of the contributing factors for the excessive generation of hyaluronan based on the fact that these cytokines are strong inducers of hyaluronan synthase-2 (HAS2) ^34^. The present report unfolded that mdig is largely responsible for the expression of inflammatory cytokines and the key component of the inflammasome, as well as most of the genes in glycan metabolism, such as Has1, Has2 and Has3 for hyaluronan generation, and GALs for O-glycosylation (Fig. 4). These effects of mdig are also attributable to the pulmonary fibrosis among some COVID-19 survivals ^22,23^. The elevated level of mdig in fibrotic human lung tissues (Fig. 6) supports this notion. Thus, over-expression of mdig, either under some chronic disease conditions or environmental exposure, possibly indicates the pathogenetic mechanism behind the severity or mortality of COVID-19. It also yields a conceivable explanation of how environmental exposure exacerbates the severity and/or mortality of COVID-19. A more complete understanding of mdig or its regulatory pathways, accordingly, could catalyze effective management strategies that will ameliorate the infectivity of SARS-CoV-2 and improve the outcomes of the COVID-19 patients.

## Materials and methods

### Cell culture

The human bronchial epithelial BEAS-2B cells were purchased from America Type Culture Collection (ATCC). BEAS-2B cells were cultured in DMEM-high glucose medium (Sigma, cat.no. D5796) supplemented with 5% FBS, 1% penicillin-streptomycin (Gibco, cat.no. 15140122), and 1% L-Glutamine (Gibco, cat.no. 25030164). Cells were maintained in a humidified incubator at 37 °C with 5% CO_2_.

### Establishment of mdig knockout (KO) cells by CRISPR-Cas9

For generation of CRISPR-Cas9 plasmids, sgRNA (5’-AATGTGTACATAACTCCCGC-3’) was selected and cloned into pSpCas9(BB)-2A-Blast vector as described ^15^. BEAS-2B cells (2.5 × 10^5^/well in 6-well plate) were transfected with CRISPR-Cas9 plasmids by Lipofectamine 2000 (Thermo Fisher Scientific) according to the manufacturer’s protocol. The transfected cells were selected with 2 *μ*g/ml Blasticidin (Thermo Fisher Scientific) until single colonies were formed. WT and mdig KO colonies were validated by measuring mdig protein expression.

### SARS-CoV2-Spike transfection

WT and mdig KO cells (3 × 10^5^/well in 6-well plate) were seeded, and grown to 70% confluence. SARS-CoV2-Spike plasmids (Addgene, cat.no. 145032) were transfected into cells using Lipofectamine 2000 (Thermo Fisher Scientific) according to the manufacturer’s protocol. Twenty-four hours after transfection, cells were subjected to western blotting.

### Western blotting

Cells were lysed in 1 × RIPA buffer (Cell Signaling), supplemented with 1mM PMSF and protease inhibitor cocktail (Thermo Fisher Scientific). Protein concentrations were determined by Pierce BCA Protein Assay Kit (Thermo Fisher Scientific). Whole cell lysates (30 *μ*g) were separated in sodium dodecyl sulfate-polyacrylamide gel electrophoresis (SDS-PAGE), and then transferred to Polyvinylidene fluoride (PVDF) membranes (Millipore). The membranes were blocked with 5% nonfat dry milk and then probed with primary antibodies at 4°C overnight. The primary antibodies include anti-mdig (Invitrogen cat.no. 39-7300), anti-SARS-CoV-2-Spike (Abcam, ab272504), anti-Furin (Abcam, ab183495), anti-TMPRSS2 (Proteintech, cat.no. 14437-1-AP), anti-Cathepsin D (Cell signaling, cat.no. 2284S), and anti-GAPDH (Cell signaling, cat.no. 2118S). After washing with TBST (3 × 10 min), the membranes were incubated with horseradish peroxidase-coupled goat anti-rabbit or -mouse IgG at room temperature for 1h and washed with TBST (3 × 10 min). Chemiluminescent Substrate (Thermo Fisher Scientific) was used to detect immunoblotting signals.

### Chromatin immunoprecipitation-sequencing (ChIP-seq)

Ten million WT or KO Beas-2B cells were fixed and subjected to ChIP with ChIP-grade antibodies against H3K9me3, H3K27me3, and H3K4me3 (Active Motif, CA). ChIP, input/control DNA libraries, NGS sequencing, and data analysis were performed as we previously described ^15^.

### RNA sequencing (RNA-seq)

Two million WT or KO BEAS-2B cells were collected for total RNA isolation, libraries preparation and NGS sequencing. The procedures and data analysis were followed as we previously reported ^15^.

### Immunohistochemistry

Pulmonary Interstitial Fibrosis tissue microarray LC561 was purchased from the USBiomax, Inc. Tissue microarray slides were processed for immunohistochemical staining for mdig as described previously ^24^. Briefly, paraffin-embedded tissue sections were deparaffinized with xylene and hydrated in a series of alcohol gradients. To quench endogenous peroxidase activity, slides were incubated with 1.5 to 3% H_2_O_2_ in PBS for 20 min at room temperature. Heat-mediated antigen retrieval was performed by boiling tissue sections in citrate buffer with pH 6.0 for 20 min in a microwave. To block nonspecific binding of immunoglobulin, slides were incubated with a solution containing 5% goat serum, 0.2% Triton X-100 in PBS for 2 h at room temperature, followed by incubation with primary antibodies against mdig (mouse anti-MINA, Invitrogen with 1:50 dilution) overnight at 4 °C. Goat anti-mouse biotinylated secondary antibodies were subsequently applied at 1:200 dilution and incubated for 2 h at room temperature. Slides were then incubated with ABC reagent (Vectastatin Elite ABC kit) for 45 min at room temperature, and the chromogen was developed with diaminobenzidine (DAB). Slides were counterstained with hematoxylin (Sigma-Aldrich, St. Louis, MO) and mounted with Entellan® (Electron Microscopy Sciences, Hatfield, PA). All incubation steps were carried out in a humidified chamber, and all washing steps were performed with 1 × PBS. Images were captured under the bright field of a Nikon Eclipse Ti-S Inverted microscope (Mager Scientific, Dexter MI, USA) and analyzed using Nikon’s NIS Elements BR 3.2 software.

### Statistics and data analysis

Expression profiles of GSE149312, GSE150392, and GSE147507 were obtained from the public NCBI-GEO database. Gene set enrichment analyses comparing RNA-seq analyses were performed using the online public database Enrichr (http://amp.pharm.mssm.edu/Enrichr/). Pathway analyses were performed by Reactome Knowledgebase (http://www.reactome.org). For Western blotting, at least three independent experiments were performed. Statistical significance for some quantitative data was determined by unpaired Student’s t-test.

## Acknowledgments

This work was partially supported by National Institute of Health grants R01 ES028263, R01 ES028335, and R01 ES031822-01A1 to FC

## Supplement figure legends

**sFig. 1.**
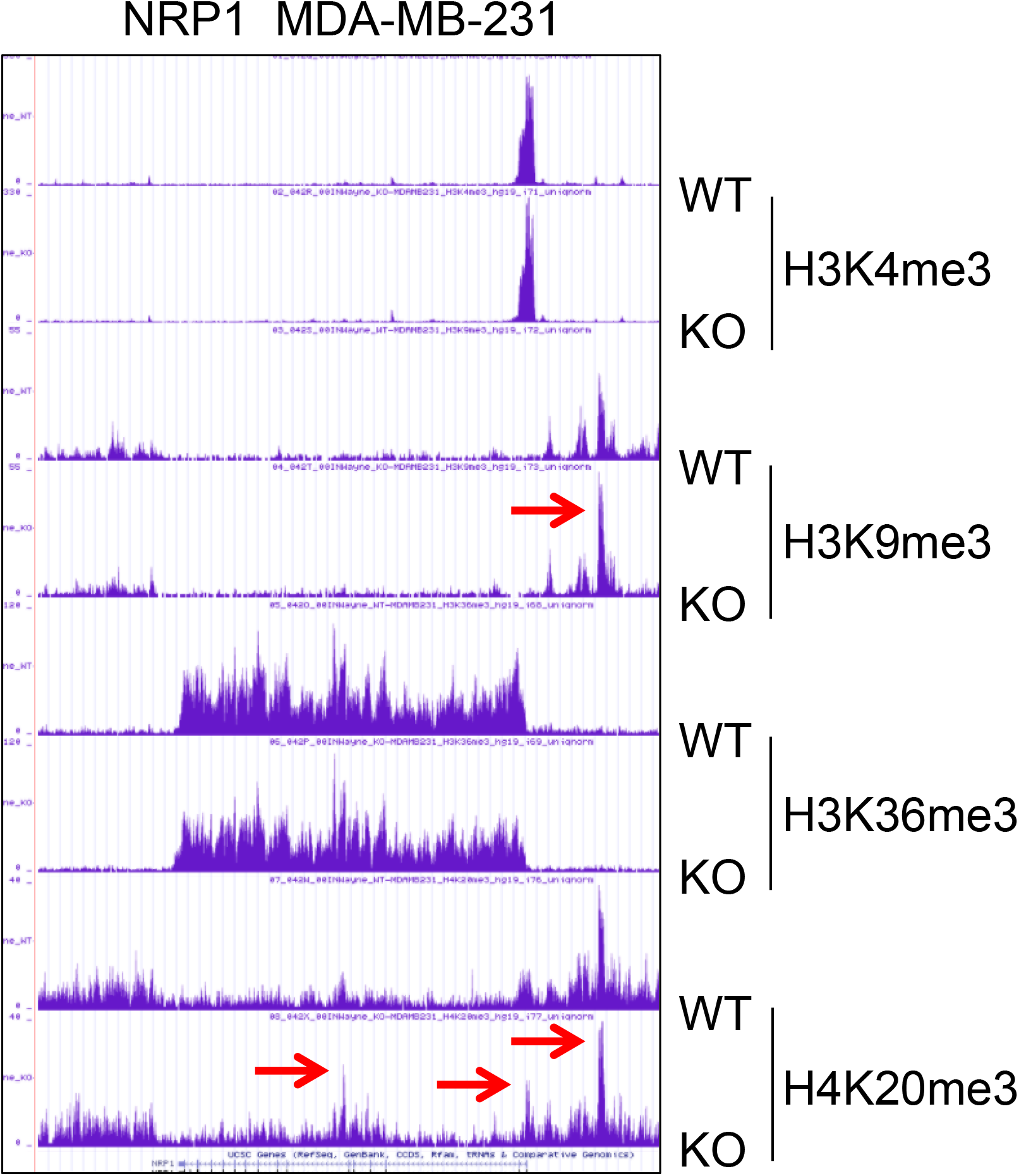
ChIP-seq of NRP1 gene in WT and mdig KO MDA-MB-231 breast cancer cell line. Red arrows point to the enhanced enrichment of H3K9me3 and H4K20me3 in mdig KO cells.

**sFig. 2.**
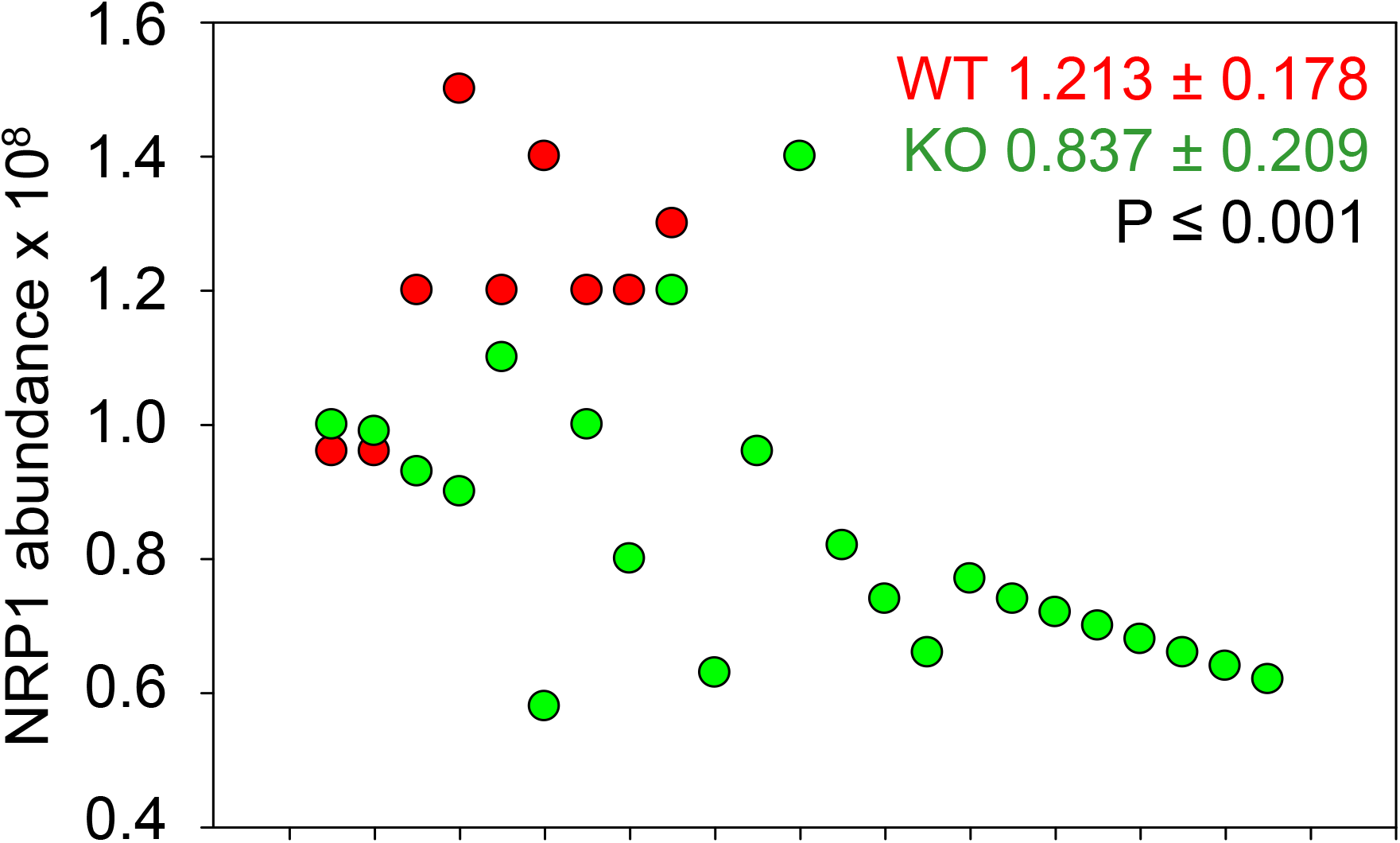
Quantitative proteomics analysis for the protein levels of NRP1 in 5 WT (red) and 11 mdig KO (green) MDA-MB-231 cell clones. Duplicated samples for each clone were subjected to quantitative proteomics analysis. One sample in each group was discarded in this analysis due to poor quality from sample preparation and processing.

**sFig. 3.**
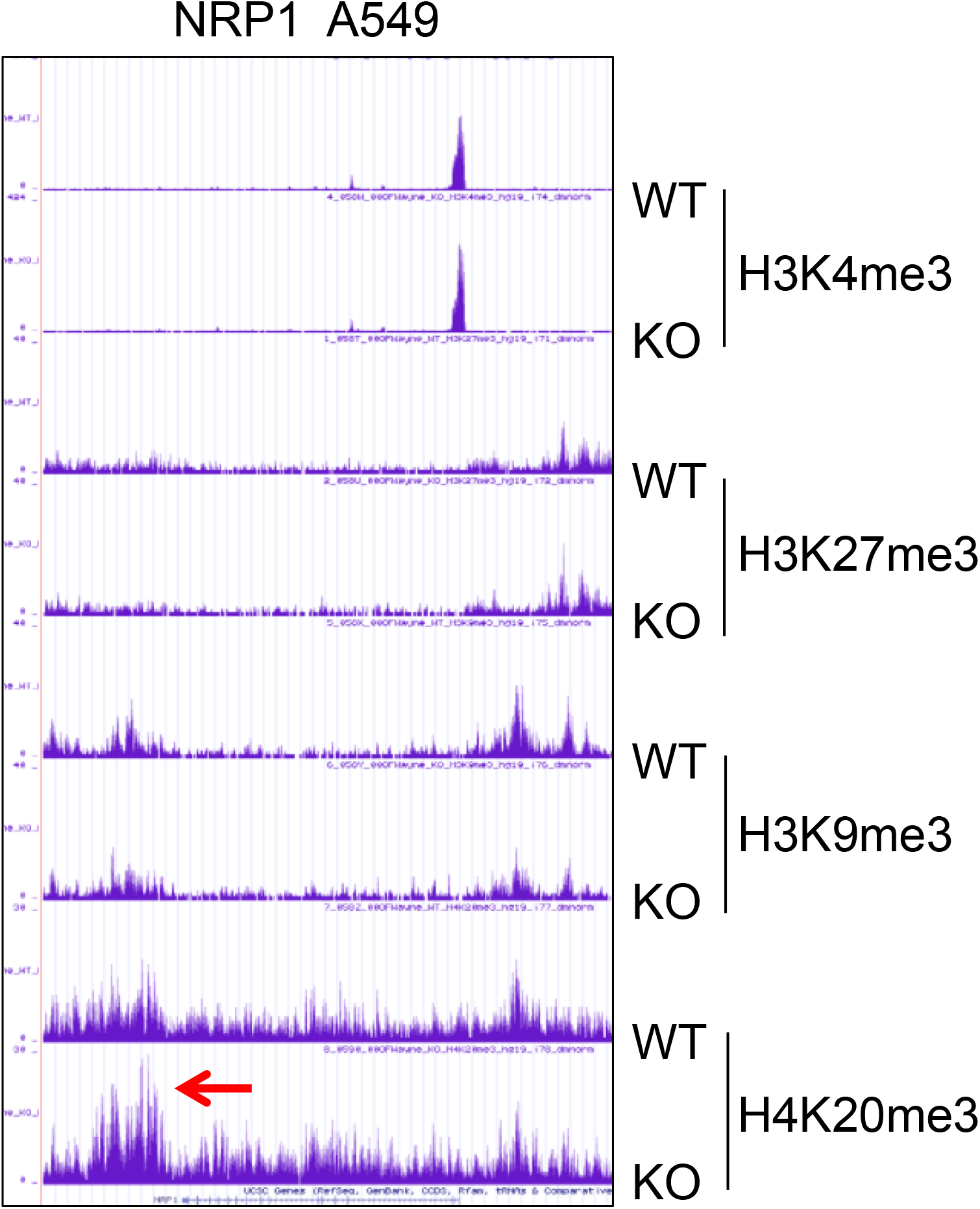
ChIP-seq of NRP1 gene in WT and mdig KO A549 cells. Red arrow points to the elevated enrichment of H4K20me3 on the NRP1 gene in mdig KO A549 cells.

